# Characterization of anaerobic bacterial populations putatively involved in the degradation of Chlorine pesticides in mangrove soils of Guadeloupe (Lesser Antilles)

**DOI:** 10.64898/2026.11.12.766228

**Authors:** E Ousselin, O. Gros

## Abstract

Due to their position at the land-sea interface and their proximity to contaminated agricultural areas, mangroves are ecosystems potentially exposed to chlordecone pollution. This study aims to describe, for the first time, the composition of bacterial communities in mangrove soils and their distribution according to chlordecone contamination.

The results revealed marked inter-site heterogeneity in bacterial communities across the three study sites, encompassing both an uncontaminated mangrove (Manche à eau; chlordecone concentrations below the detection limit, <5 µg kg^−1^) and contaminated mangroves (Sarcelle and Petite rivière de Goyave), where soil chlordecone concentrations ranged from <5 to 118.8 µg kg^−1^ along the coastal-to-agricultural transects. Between 60 and 90% of the detected taxa belong to groups involved in the degradation of aromatic compounds, while 5 to 15% correspond to genera associated with nitrification processes. This diversity does not appear to be directly correlated with the presence of chlordecone, suggesting the predominant influence of other environmental factors. Nevertheless, two bacterial genera of interest were identified, *Dehalogenimonas* and *Desulfomonile*, known for their involvement in the degradation of aromatic compounds according to the FAPROTAX analysis.

Furthermore, functional analysis based on the KEGG database suggests the presence of vitamin B12 biosynthesis pathways in *Dehalogenimonas* and *Desulfomonile* (found in coastal areas), as well as in *Burkholderia* (found in agricultural soils). These results suggest the existence of metabolic potential to intervene in both the chemical and biological processes of chlordecone degradation.

Finally, mangroves appear to have purification potential favorable to chlordecone degradation, due to the presence of *Dehalogenimonas* and several other bacterial genera capable of participating in the chemical and/or biological degradation processes of chlordecone.

## I. Introduction

In the late 1960s, the weevil *Cosmopolites sordidus*, a crop pest, caused significant losses in banana plantations in the French islands (Guadeloupe and Martinique) in Caribbean (Dromard et al., 2018). To combat this invasion, a pesticide called chlordecone was used. In Guadeloupe, its application was mainly concentrated in Basse-Terre due to the prevalence of banana plantations in this region; consequently, our sampling strategy was focused there. Discovered in 1951 and patented in 1952 by Allied Chemical Corporation, chlordecone (C_10_Cl_10_O) is an organochlorine compound with a unique chemical structure, called a bishomocubane cage, which gives it chemical stability. It was originally registered and marketed under the trade name Kepone® (Wilson, 2023) before being officially designated as chlordecone. Chlordecone has been classified as a persistent organic pollutant (POP) by the Stockholm Convention because of its high hydrophobicity, which allows it to adsorb more easily to soil components (Cabidoche et al., 2009). The persistence of chlordecone depends on the type of soil: approximately 70 years in nitisols versus nearly 400 years in andosols. In 2001, surveys conducted by the French Ministry of Health revealed the presence of chlordecone in soil, surface water, groundwater, and various agricultural products, particularly root vegetables (Dromard et al., 2018). Indeed, the high concentration of chlordecone in soil, estimated between 0.2 and 37.4 mg.kg^−1^ (Cabidoche et al., 2009), has led to long-term contamination of groundwater and intertidal areas, such as estuaries and mangroves, through rainwater runoff (Coat et al., 2006; Merlin et al., 2016). As a result, fishing bans have been imposed in certain areas due to the bioaccumulative nature of chlordecone in many aquatic organisms (Dromard et al., 2018). In terms of health, chronic exposure to chlordecone has been linked to an increased risk of prostate cancer and fertility problems in the French West Indies (Sanchez et al., 2025).

Long considered non-degradable, chlordecone has nevertheless been found to be capable of “transforming” into degradation products, observed both in the laboratory and *in situ* in the soils of the French islands in the Lesser Antilles (Lomheim et al., 2021). This transformation capacity paves the way for in-depth research into possible remediation strategies aimed at exploiting these processes to mitigate this persistent pollution. Two main hypotheses have been proposed to explain the degradation of chlordecone. The first corresponds to a biological pathway involving bacteria of the genera *Citrobacter*, *Desulfovibrio*, and *Pseudomonas* (Figure 1), capable of catalyzing dehalogenation or reductive sulfidation reactions (Barbance et al., 2020; Della-Negra et al., 2021). The second hypothesis is based on chemical degradation in the presence of reducing agents and cofactors such as vitamin B_12_ (Figure 1), also known as cobalamin (Ranguin et al., 2017). Vitamin B_12_ plays an important role as an electron carrier in the reduction of halogenated compounds in the environment (Guo and Chen, 2018). It consists of three parts: a central corrin ring, an upper axial ligand, and a lower ligand (Guo and Chen, 2018). In humans and other organisms, vitamin B_12_ plays a role in various metabolic processes. In prokaryotes, cobalamin supports the fermentation of small molecules and catalyzes molecular rearrangement. Cobalamin acts as a cofactor for a key enzyme, reductive dehalogenase (Payne et al., 2015). This enzyme is involved in numerous pathways for the transformation and degradation of halogenated compounds, catalyzed by various anaerobic bacteria.

**Figure 1:**
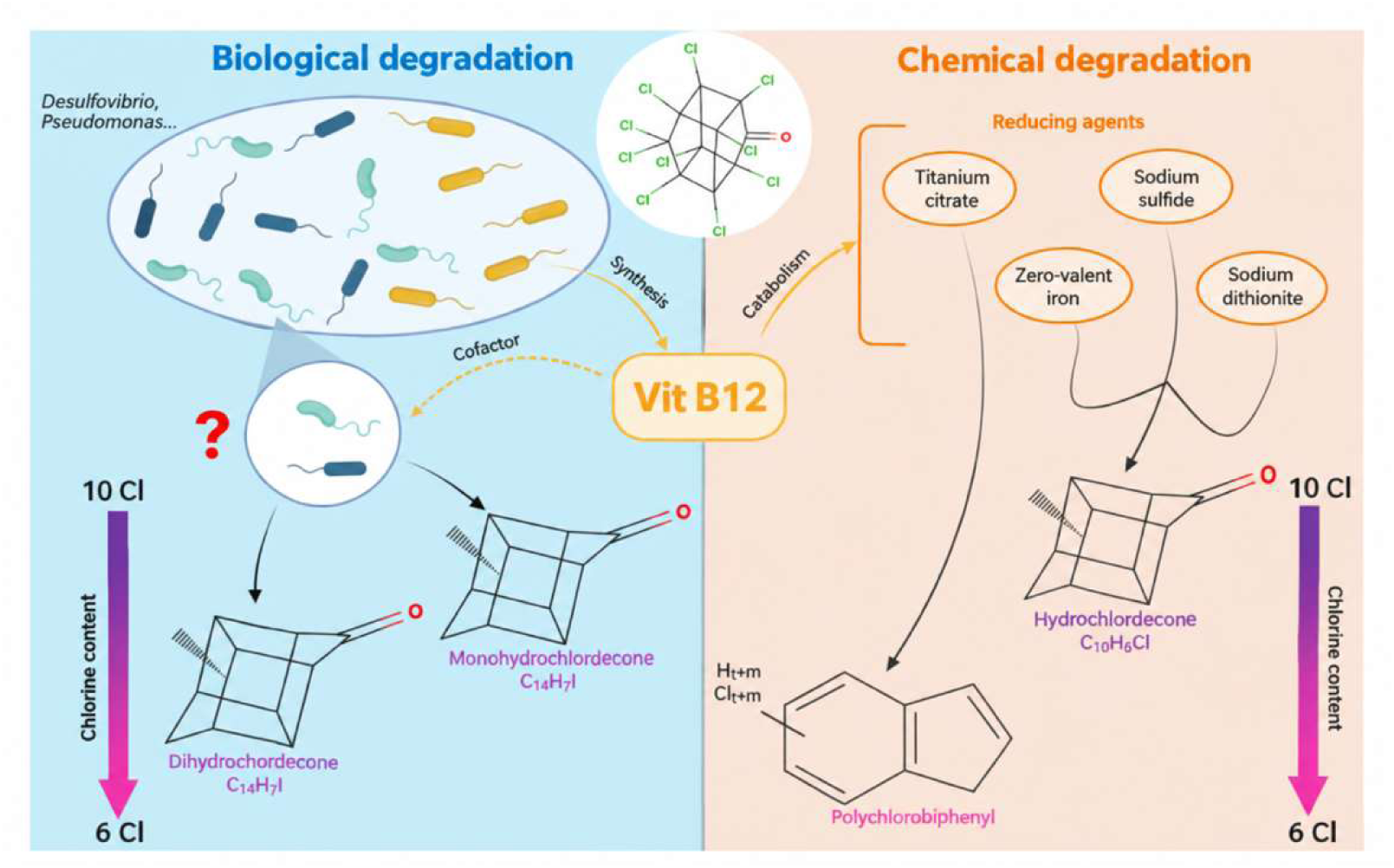
Chlordecone Degradation: The Key Role of Vitamin B12 in Biotic and Abiotic Pathways. Chlordecone is shown at the center of the figure, with its two main degradation pathways: biological (in blue) and chemical (in orange). Vitamin B12, positioned centrally, acts as a cofactor. On the biological side (in blue), anaerobic bacteria participate in reductive dehalogenation reactions, leading to the formation of dihydrochlordecone and monochlordecone. On the chemical side (in orange), various reducing agents associated with vitamin B12 induce the transformation of the molecule into polychloroindene and hydrochlordecone. Diagram created using BioRender, based on Chaussonnerie et al. (2016) and Chevallier et al. (2019).

Thus, combining the biological production of this cofactor with the chemical degradation of chlordecone by reducing agents could be a viable strategy (Chaussonnerie et al., 2016; PL Saaidi, personal communication). For this reaction to occur (Figure 1), certain conditions are required, such as the presence of a carbon source acting as an electron donor and an anoxic environment (Lomheim et al., 2021). These conditions are met in the mangroves of Guadeloupe, making this ecosystem an ideal model for bacterial processes occurring in anaerobic conditions (Crémière et al., 2017; Sebilo et al., 2026).

Mangroves, tropical ecosystems of Guadeloupe’s marine waters, line the coastline creating a unique environment characterized by singular ecological challenges (Gontharet et al., 2017). Indeed, the sediments of coastal mangroves experience salinity levels of around 35 g L^−1^, marked anoxic conditions, and a substrate in perpetual sedimentation (Gontharet et al., 2017). In contrast, mangrove soils (terrestrial mangroves) are subject to decreasing salinity from the coastline to agricultural fields, gradually decreasing from 25 g L^−1^ to less than 1 g L^−1^ in the agricultural fields (Devarieux, 2025). Due to their location at the interface between land and sea, mangroves are potentially exposed to contamination by chlordecone. In the mangroves of southern Basse-Terre, located in areas where this pesticide has been used, pollution results mainly from runoff and leaching of rainwater. This phenomenon carries chlordecone from agricultural soils to these coastal ecosystems (Cattan et al., 2019). Monitoring stations located in the south-western part of Basse-Terre show the highest concentrations of chlordecone in fresh water, with several sites exceeding 1 µg/L and most values ranging between 0.1 and 1 µg L^−1^. These concentrations are markedly higher than those observed in the northern part of the island, where contamination is often not detected or remains below 0.01 µg L^−1^ (Rochette et al., 2017).

Furthermore, mangroves could promote the degradation of chlordecone due to their very low redox potential and their putative ability to store pollutants in their sediments (Cremière et al., 2017; Lavermann et al., 2023). This purifying role has also been demonstrated in other contexts: for example, anaerobic degradation of sulfamethoxazole has been observed in mangrove sediments in Taiwan (Yang et al., 2025). Bacterial populations and associated functions will evolve according to salinity gradient from the marine fringe to the swamp forest, with significant ecological implications. For example, bacterial communities in marine mangroves exhibited higher diversity as revealed by 16S rRNA gene sequencing analysis, with the presence of diverse bacterial phyla including Proteobacteria, Actinobacteria, and Bacteroidetes (Gómez-Acata et al. 2023, Jamon-haon et al. 2025; Hervé et al., 2026). Unfortunately, these studies did not focus on bacteria putatively involved in the bioremediation of chlorine pesticides

In this context, this study aims to identify bacteria potentially involved in the degradation process of chlordecone in mangroves, an ecosystem that has been little studied despite its interesting biotic and abiotic characteristics. A metabarcoding approach will be used to analyze the bacterial diversity of these environments and to compare the taxonomic composition and predictive metabolic functions (FAPROTAX) between different contaminated areas. We hypothesize that mangroves could harbour more diverse bacterial communities that are more specialized in biogeochemical cycles, including bacteria of interest for the bioremediation of chlordecone.

## II. Materials and Methods

### 1. Sampling

Soil sampling was carried out at Manche à Eau (16°16.608′N, 61°33.425′W), along a transect extending from the seaside, through the mangrove, to the fields (cultivated or not). This transect was used to collect representative samples from different zones: seaside, middle of mangrove, back mangrove and fields adjacent to mangrove (agricultural area). For each zone, samples were taken to a depth of at least 5 cm using an auger. The Manche à Eau site served as a negative control (Figure 1B) because analyses have confirmed the absence of chlordecone (detection threshold < 5 μg/kg, this study) and low trace metal concentrations (Colletin et al., 2026).

The same strategy was applied at the Sarcelle site (16°09.591′N, 61°34.921′W) and at the mouth of the Petite rivière de Goyave (16°08.059′N, 61°33.984′W) (Figure 2). Chlordecone concentrations were determined in soil samples collected from the Sarcelle and Petite Rivière de Goyave transects. In addition, soil samples were collected for bacterial community analysis by 16S rRNA gene metabarcoding. For each sample, approximately 0.1 g of soil was subsampled and stored at −80°C until DNA extraction and subsequent molecular analyses.

**Figure 2:**
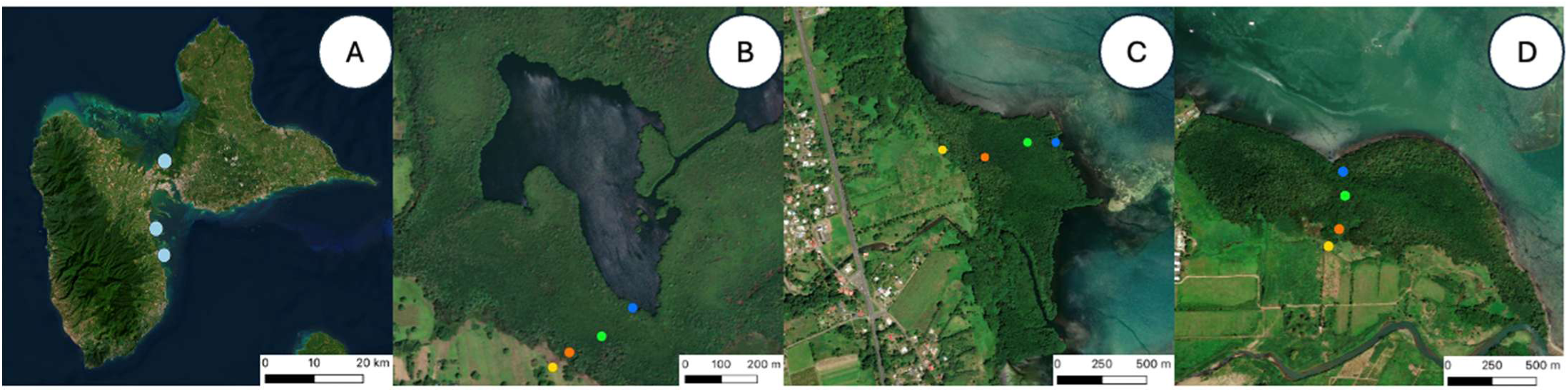
Location of sampling sites in the mangroves of Guadeloupe. (A) Overview of the three mangrove areas studied in (A) Guadeloupe: (B) Manche-à-Eau, (C) Sarcelle, (D) Petite Rivière de Goyave. For each site, a four-point transect was carried out: Seaside (blue), Middle of mangrove (green), Back mangrove (orange), Field adjacent to mangrove (yellow). Maps created using QGIS 3.34 based on ESRI satellite imagery (ESRI Satellite).

### 2. Chlordecone measurement from soils

Chlordecone concentrations were determined by the Drôme Laboratory (Terrana Drôme). Samples were collected, stored at −20°C until analysis, and quantified using high-performance liquid chromatography coupled with mass spectrometry (HPLC–MS).

### 3. Sequencing of 16S rRNA genes

Bulk DNA was extracted from each slide using the FastDNA™ SPIN Kit for Soil (MPBio). The quantity and quality of DNA were measured using NanoDrop (Thermo Scientific). The extracted genomic DNA was stored at −20 °C until PCR amplification. The conditions for PCR amplifications were defined as follows: initial denaturation at 95 °C for 3 min; 29 cycles at 95 °C for 45 s; annealed at 50 °C for 45 s; extension at 72 °C for 1 min 30 s; and a final extension of 5 min at 72 °C. The V3 and V4 regions of the microbial 16s rRNA were amplified using two universal PCR primers (Bac357F [α-CCTACGGGAGGCAGCAG] and Bac926R [β-CCGTCAATTCMTTTRAGT]). The PCR products were performed in triplicates and sent to the company Microsynth (9436 Balgach, Switzerland) for high-throughput sequencing steps and the construction of the OTU abundance tables.

#### 3.1. Amplicon data analysis

The produced paired-end reads were subject to de-multiplexing and trimming of Illumina adaptor residuals using Illumina’s bcl2fastq software version v2.20.0.422. The quality of the reads was checked with the software FastQC version 0.11.8 and sequencing reads that fell below an average Q-score of 24 or had any uncalled bases (N) were removed from further analysis. The locus specific V34 primers were trimmed from the sequencing reads with the software cutadapt v3.2. Trimmed forward and reverse reads of each paired-end read were merged to in-silico reform the sequenced molecule considering a minimum overlap of 15 bases using the software USEARCH version 11.0.667. Merged reads that contained ambiguous bases or were outliers regarding the expected amplicon size distribution were also discarded. Samples that resulted in less than 5000 merged reads were discarded, to not distort the statistical analysis. The surviving reads were denoised to form zero-radius OTUs (zOTUs) discarding singletons and chimeras in the process. The resulting zOTU abundance table was then filtered for possible barcode bleed-in contaminations using the UNCROSS algorithm. The zOTU sequences were compared to the reference sequences of the NCBI RefSeq Targeted Loci database.

### 4. Statistical analyses, functional group prediction

Shannon and Simpson alpha diversity indices were calculated on R (v2023.09.1+494) using the Vegan package. Several statistical tests were used to process the data.

The prediction of functional bacterial groups was performed by cross-referencing OTU data from 16S rDNA sequences with the FAPROTAX database (Louca et al., 2016). Each OTU is assigned to groups involved in metabolic functions related to different biogeochemical cycles and the degradation of organic matter (Yang et al., 2022). The relative abundance of OTUs involved in these functions was calculated for the three sites studied.

An analysis of the metabolic pathways involved in vitamin B_12_ biosynthesis was performed for the most abundant bacterial genera in samples from the three sites, using the KEGG platform. This database brings together various metabolic maps representing current knowledge on molecular interaction networks and biological reactions.

A search was also conducted for the presence of the enzyme of interest, reductive dehalogenase, which is involved in biological degradation reactions.

## III. Results

### I. Chlordecone measurement from soils

Soil analyses performed by the TERRANA laboratory revealed contrasting chlordecone contamination profiles among the three study sites. At Manche-à-Eau, (chlordecone concentrations were consistently below the analytical limit of detection (<5 µg kg^−1^) throughout the transect. In contrast, measurable chlordecone concentrations were detected at Sarcelle and Petite rivière de Goyave sites, displaying an overall increasing contamination gradient from the seaside to the fields adjacent to mangrove. At Sarcelle site, chlordecone concentrations ranged from <5 µg kg^−1^ at the seaside to 118.8 µg kg^−1^ in the agricultural fields, with intermediate concentrations of 42.4 and 26.8 µg kg^−1^ in the middle of mangrove and back-mangrove, respectively. At Petite rivière de Goyave, concentrations increased from 9.6 µg kg^−1^ at the seaside to 41.3 µg kg^−1^ in the agricultural fields, although a slight decrease was observed between the mangrove (32.1 µg kg^−1^) and the back-mangrove (27.9 µg kg^−1^).

### II. Taxonomic comparition of soil samples explored by metabarcoding analysis of the bacterial gene encoding 16S rRNA

Chlordecone analyses confirming high levels of contamination in our study areas, Petite rivière de Goyave and Sarcelle. Taxonomic analysis of soil samples revealed 26 bacterial phyla, comprising 59 classes, 107 orders, 183 families, 305 genera, and a total of 388 species present in the samples. In the next stage of this work, bacterial composition was studied at the genus level.

The relative abundance profile (%) shows marked population heterogeneity for the three sites sampled: Manche à Eau, Sarcelle, and Petite rivière de Goyave. Analysis of the relative abundances of bacterial genera reveals three levels of structure.

The first dimension of the analysis shows a bacterial structure specific to the sampled sites, marked both by diversity and by the dominance of certain genera (Figure 3). Firstly, much greater diversity is observed at the two sites exposed to chlordecone (Sarcelle and Petite rivière de Goyave) and much lower diversity at Manche à Eau (not impacted by chlordecone). The Manche à Eau site has a high diversity of bacterial genera, with a relative dominance of *Dehalogenimonas*, *Desulfatiglans*, and *Desulfatalea*. The Sarcelle site has a predominance of the bacterial genera *Desulfomonile*, *Gemmatirosa*, *Sulfuritortus*, and *Thiohalophilus*. Finally, the Petite rivière de Goyave site is characterized by a high abundance of *Thermodesulfovibrio*, *Thiohalophilus*, and *Desulfomonile*.

**Figure 3:**
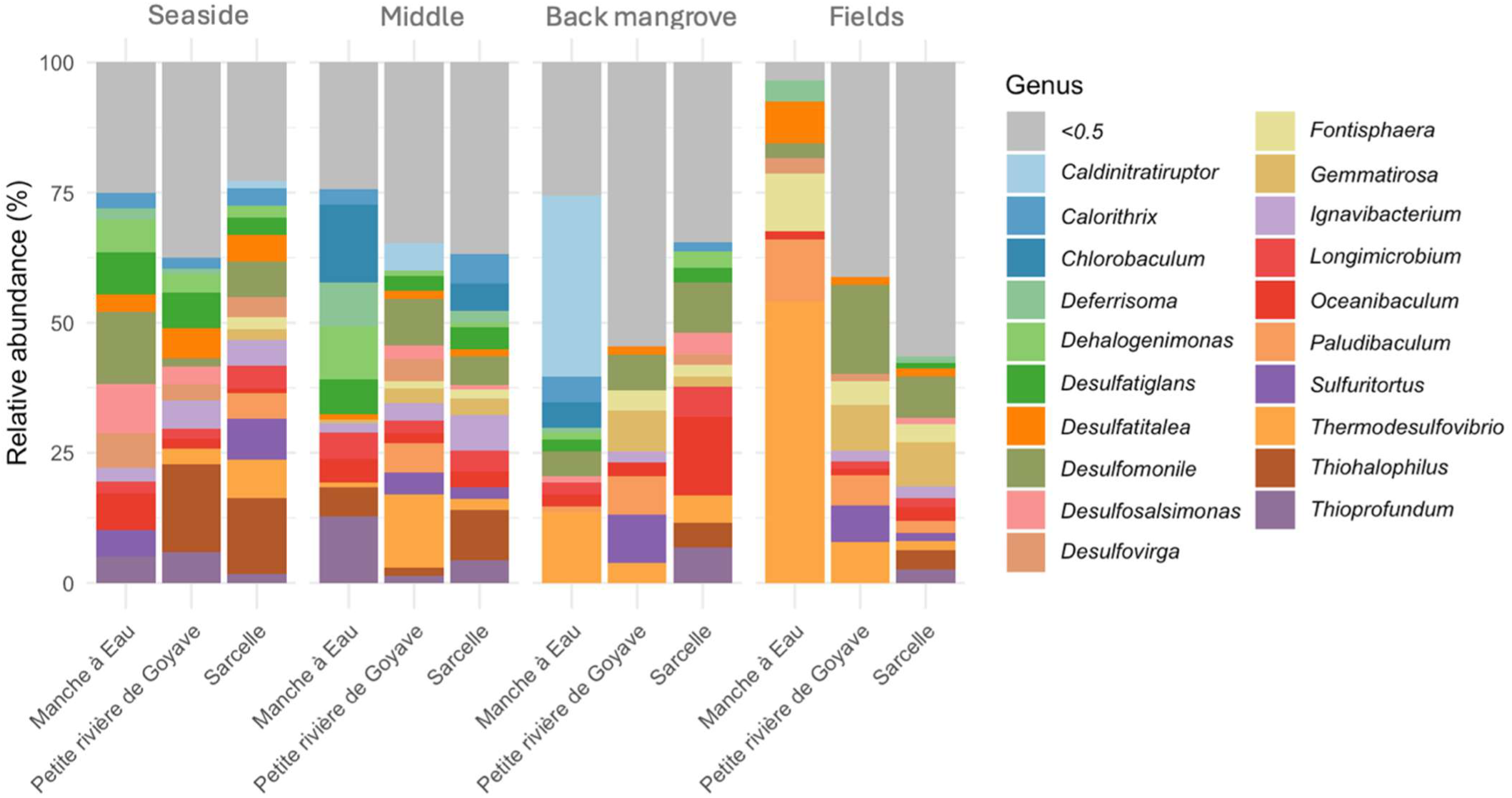
Metabarcoding profile of bacterial genera observed at the three sites. The relative abundance (%) of all samples from the transect was calculated.

At the intra-site level, bacterial composition varies between samples, with greater diversity observed at certain sites (Figure 3). The Manche à Eau site shows variability in bacterial genera along the entire transect. Indeed, samples from the seaside show a high abundance of the genera *Dehalogenimonas*, *Desulfomonile*, and *Desulfosalsimonas*, but these are not found along the transect. Soils in the middle of the mangrove are characterized by a predominance of the genera *Chlorobaculum* and *Dehalogenimonas*. Finally, towards the rear mangrove and agricultural fields, the soils are marked by a majority of bacteria belonging to the genera *Burkholderia*, *Desulfatitalea*, and *Thermodesulfovibrio*.

In Sarcelle, the profiles are quite similar, but with varying abundances along the transect. The samples taken at the seaside are characterized by a predominance of the genera *Desulfomonile*, *Thiohalophilus*, and *Thermodesulfovibrio*. The samples taken from the “middle” of the mangrove have profiles similar to those taken at the seaside, but with a predominance of the genera *Ignavibacterium*, *Sulfuritortus*, and *Thiohalophilus*. The soil sample taken from the back mangrove still has a profile similar to that of the seaside, but with an abundance of the bacterial genera *Desulfomonile*, *Ferruginivarius*, and *Oceanibaculum*. Finally, soil samples taken from fields adjacent to mangrove show a predominance of the genera *Desulfomonile* (found along the transect), *Gemmatirosa*, and *Sulfurovum*.

The Petite rivière de Goyave site is characterized by heterogeneity in bacterial genera along the transect, although samples from the coast appear to be relatively less diverse than others. The same bacterial genera, such as *Desulfomonile*, *Desulfatitalea*, *Thiohalophilus*, and *Thermodesulfovibrio*, are found throughout the transect. As we move inland, away from the coast, an increase in the relative abundance of *Desulfomonile*, *Gemmatirosa*, and *Sulfuritortus* is observed, particularly in areas close to agricultural fields.

The results show that each site studied has its own unique structure. Each site has a distinct bacterial composition that reflects the strong influence of environmental conditions. The Manche à Eau site, used as a control site unaffected by chlordecone pollution, shows much lower diversity along the transect than the two contaminated sites (Sarcelle and Petite rivière de Goyave). These sites affected by chlordecone contamination are characterized by the predominance of genera such as *Thiohalophilus* and *Thermodesulfovibrio*. This contrast suggests that chlordecone could positively influence species richness by promoting more diverse and specialized bacterial communities.

At the intra-site level, the spatial gradient observed along the transects from the seaside to the fields adjacent to mangrove shows dominant genera that vary gradually depending on the transect sampling area. For example, the genera *Desulfomonile* and *Sulfuritortus* tend to be more abundant in samples collected from fields near mangroves.

Finally, the proportion of taxa with a relative abundance <0.5 varies greatly from one site to another, which could reflect differences in terms of species richness or alpha diversity. Thus, the overall heterogeneity observed between sites, along transects, and as a function of depth reflects the complexity of bacterial dynamics in the mangroves of Guadeloupe. In order to better understand the structure and distribution of these bacterial communities, a diversity analysis is needed to complement the initial observations of the Metabarcoding profile.

### 2. Analysis of alpha diversity in soil samples (between each site)

The richness observed in OTU(S) differed considerably between areas contaminated and uncontaminated by Chlordecone.

Firstly, it can be seen that the Petite rivière de Goyave site appears to have the greatest diversity in terms of species (Figure 4a). Conversely, the site not contaminated with chlordecone, corresponding to Manche à Eau, appears to have lower diversity. The differences in species diversity between the three sites are statistically significant, with a p-value of 0.001623 according to the ANOVA test. These analyses confirm our interpretations of the previous metabarcoding profile (Figure 3).

**Figure 4:**
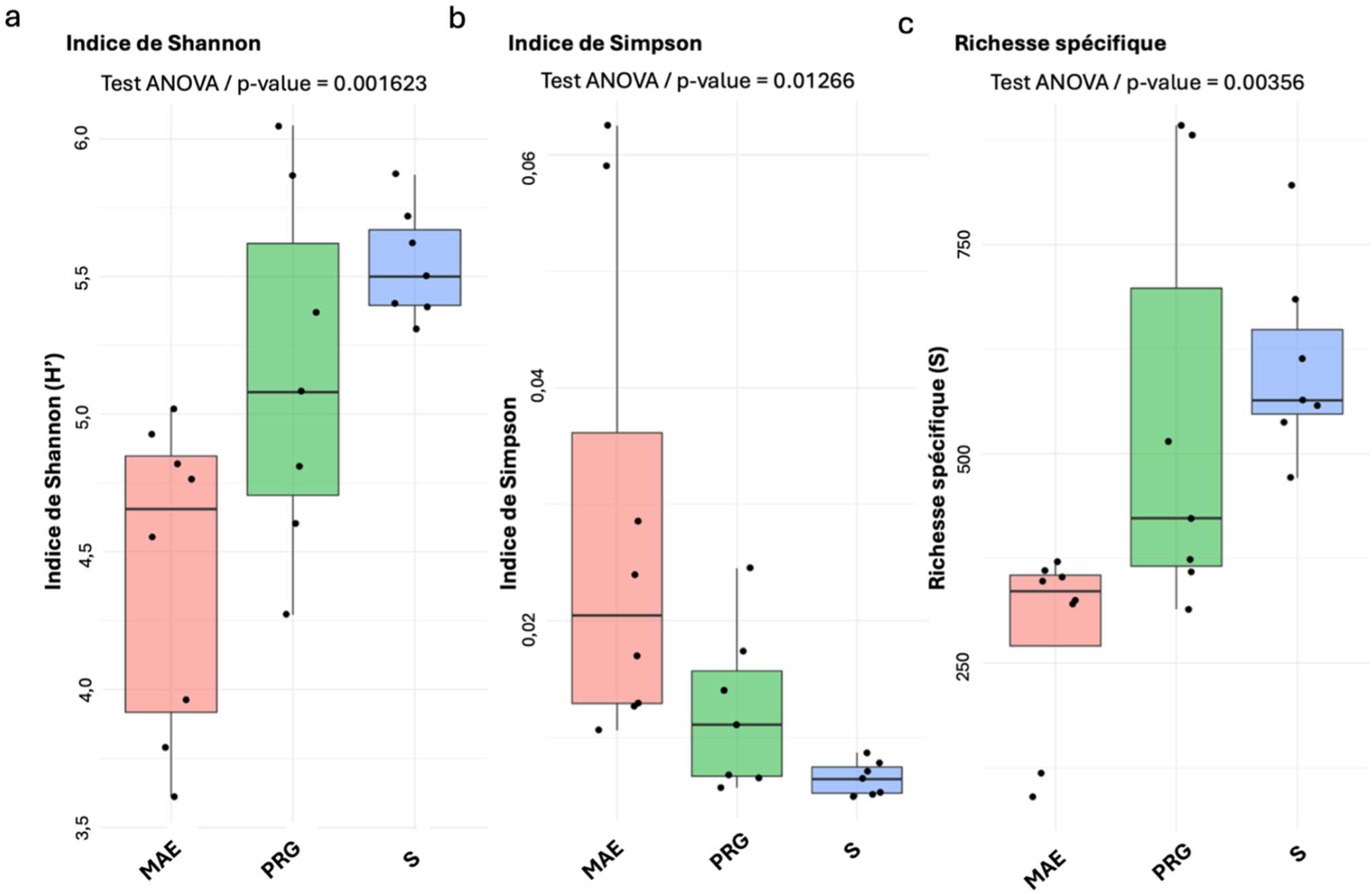
Comparison of indices associated with the study of alpha diversity of different samples from the sites. Shannon and Simpson biodiversity indices. Species richness corresponding to the total number of species in a sample. Each colour is associated with a site: red (Manche à Eau or MAE), green (Petite rivière de Goyave or PRG), and blue (Sarcelle or S). An analysis of variance (ANOVA) was performed to assess the significance of the differences observed between the sites studied. (n=8) for each boxplot.

We observed that Manche à Eau has a Simpson index close to 1, indicating that bacterial genera are numerous and distributed in comparable proportions (Figure 4b). By comparison, Sarcelle and Petite rivière de Goyave sites have a significantly lower diversity index between 0.00 and 0.02 (ANOVA, p-value=0.0126), reflecting higher species diversity with a dominance of certain genera.

Finally, the specific richness observed shows that Manche à Eau has low richness compared to Petite rivière de Goyave and Sarcelle, which appear to be richer in bacterial genera (Figure 4c). It should be noted that for the Petite rivière de Goyave site, richness appears to be more stable and moderate, while for the Sarcelle site, richness appears to be more variable, given the error bar and the extent of the boxplots.

Thus, for each site, significant differences in diversity are observed depending on the presence or absence of chlordecone. The Manche à Eau site appears to be the least rich and least diverse overall, compared to the two contaminated sites, Sarcelle and Petite rivière de Goyave. Sarcelle has a higher species richness, accompanied by a high degree of population heterogeneity, suggesting a complex ecological dynamic. Finally, Petite rivière de Goyave combines relatively stable diversity and species richness, which could indicate a more balanced ecosystem that is potentially better adapted to contamination.

### 3. FAPROTAX analysis: relative abundance profile of OTUs detected from predictive metabolic pathways

FAPROTAX analysis was used to predict the functional groups present in the soil samples collected and to evaluate the different metabolic activities present at the three sites. During this analysis, for the “seaside” sample from the Sarcelle site, no OTU was associated with a physiological function. Firstly, it can be observed that across all sites, the respiration of sulfur compounds is a dominant process with fluctuating relative abundance (Figure 5). In addition, other metabolic processes such as nitrification and denitrification are present, but in lower proportions compared to sulfate respiration (Figure 5). At the Sarcelle and Petite rivière de Goyave sites, these two metabolic processes appear to be much more important compared to Manche à Eau, suggesting an abundance of bacterial genera associated with these metabolic functions in contaminated areas.

**Figure 5:**
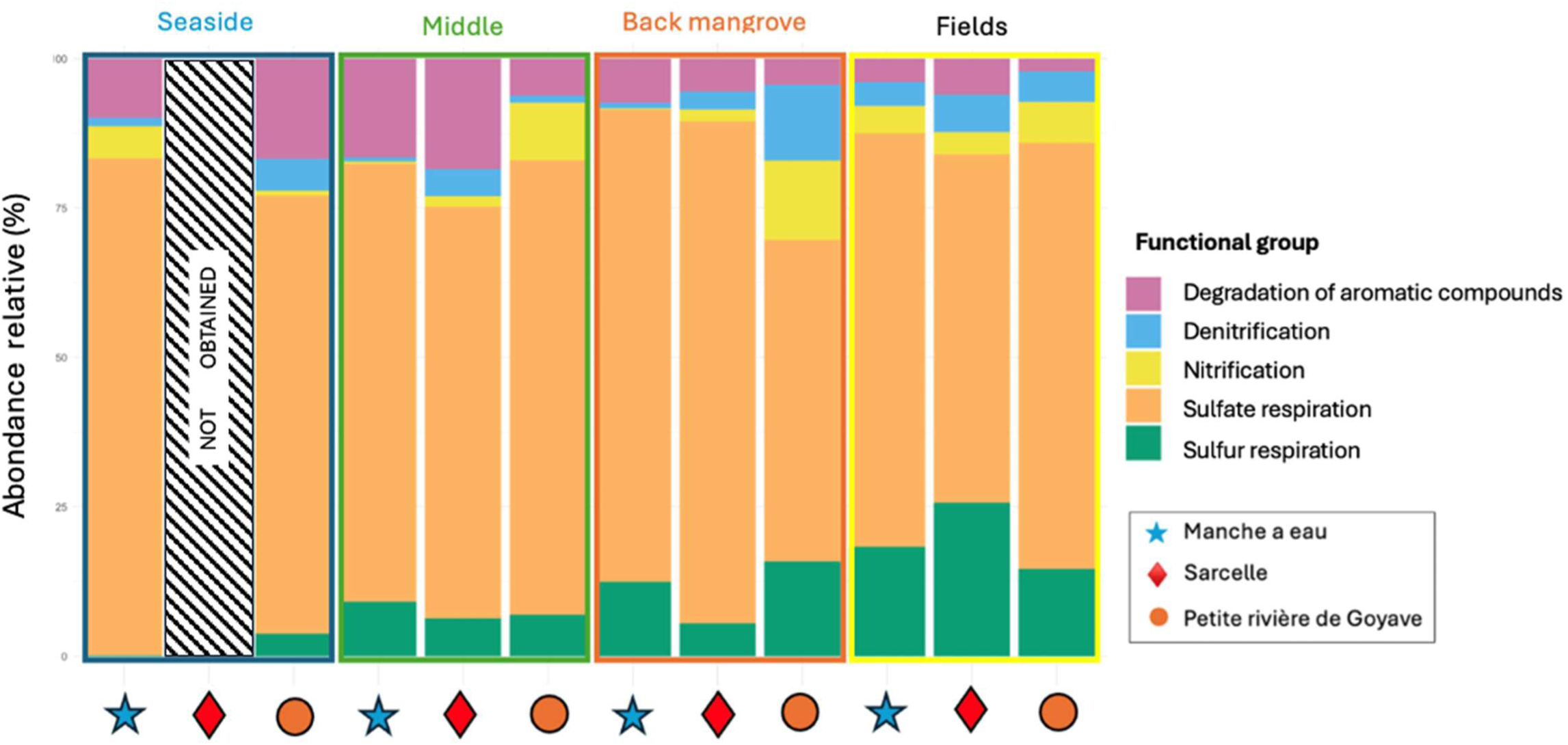
Profile of the relative abundance of functional groups present at the three sites studied.

On an inter-site scale, there is a functional transition between sulfur respiration and sulfate respiration (Figure 5). Sulfate respiration tends to decrease along the transect from the seaside to the fields. These results seem to suggest the establishment of metabolic functions along the transect requiring certain environmental factors (anoxia, humidity, sulfur/sulfate availability). At all three sites, there is a function of aromatic compound degradation, especially for samples taken at the seaside and in the mangrove environment (Figure 5).

Other processes, such as denitrification and nitrification, appear to increase along the transects (Figure 5). Indeed, these functions are dominant in the seaside, mangrove forest (middle of mangrove and back mangrove) and fields samples for all three sites. However, the putative groups with denitrification and nitrification functions are more abundant in Sarcelle and Petite rivière de Goyave.

Finally, FAPROTAX analysis made it possible to visualize the presence, or absence, of functional groups associated with metabolic functions of interest. The presence of the “aromatic compound degradation” function at the seaside and in the middle of mangrove of contaminated areas suggests that ecological specialization exists.

It could be that certain bacterial genera have adapted locally in response to complex environmental conditions. On the other hand, functions such as the respiration of sulfur and sulfate compounds confirmed the presence of bacterial genera associated with sulfate-reduction and sulfur-oxidation in mangrove soils. With regard to nitrogen, nitrification and denitrification are metabolic processes found mainly in Sarcelle and Petite rivière de Goyave. Thus, the bacterial groups performing these metabolic activities are more prevalent at these two sites. In conclusion, the spatial distribution of bacterial communities along the transect appears to be closely linked to their metabolic functions.

### 4. Analysis of the metabolic pathways of the most abundant bacterial genera

The five most abundant bacterial genera in the samples were selected for metabolic pathway analysis (KEGG). The aim of this analysis was to determine whether the bacterial genera possessed a vitamin B_12_ biosynthesis pathway and to verify the presence of a gene encoding a reducing dehalogenase.

The KEGG analysis identified four bacterial genera with a complete cobalamin biosynthesis pathway, namely *Dehalogenimonas* (Manche à Eau and Petite Rivière de Goyave), *Ilumatobacter* (Sarcelle), *Burkholderia* (Manche à Eau), and *Thermodesulfovibrio* (Petite Rivière de Goyave). The pathways involved include KEGG modules [M00122] and [M00123], corresponding to the anaerobic/aerobic synthesis of vitamin B_12_. These results suggest that these genera could contribute to the availability of vitamin B_12_ at the sites studied. In addition, among the five bacterial genera sampled, the genus *Dehalogenimonas* was found to possess a gene encoding a dehalogenase reductase. This characteristic makes it a potential candidate for the bioremediation of contaminated soils.

These results highlight the metabolic potential of bacterial genera such as *Burkholderia* and *Thermodesulfovibrio* to play a role in vitamin B_12_ availability in Manche à Eau and Petite Rivière de Goyave sites (contaminated or not). Furthermore, the predicted functional potential for vitamin B_12_ biosynthesis, together with the presence of taxa potentially associated with the degradation of chlorinated compounds, such as *Dehalogenimonas*, suggests the potential for a biological chlordecone degradation pathway at the Manche à Eau and Petite Rivière de Goyave sites. These observations pave the way for hypotheses on the use of mangrove bacterial populations in the remediation of chlordecone-contaminated soils.

## IV. Discussion

Metabarcoding analysis of mangrove soils from sites contaminated or uncontaminated by chlordecone revealed significant inter-site heterogeneity. These observations reflect a bacterial community structure influenced by local environmental parameters. Like the mangroves of Guadeloupe (Hervé et al., 2026), the mangroves of China (Yao et al., 2025) have been extensively studied for their ecological and physicochemical complexities at intervals of a few kilometres. Each mangrove area harbors specific bacterial communities adapted to environmental conditions, depending on whether they are terrestrial or marine mangroves. Bacterial communities in mangroves appear to be structured by abiotic gradients such as salinity, oxygenation, and substrate type (Alongi et al., 1993).

Heterogeneity manifests itself at several levels. First, alpha diversity indices (Shannon and Simpson) show higher microbial diversity in contaminated sites (Petite Rivière de Goyave and Sarcelle) compared to the control site (Manche à Eau). This observation could reflect a phenomenon of adaptation and/or ecological selection, whereby bacterial communities have evolved to tolerate chronic contamination by chlordecone. This type of adaptation has already been observed in agricultural environments (Brandt et al., 2009), where certain bacteria have acquired genes encoding enzymes capable of degrading toxic compounds. These enzymes allow them to use toxic compounds as a source of carbon, thus promoting their persistence in the environment. The persistence of these bacteria therefore allows for an increase in microbial diversity through the selection of tolerant taxa (Brandt et al., 2009).

The structure of communities observed according to site dimensions may be the result of several factors. Factors such as site topography, particularly soil type, could influence bacterial distribution. According to previous analyses, contaminated soils (Petite Rivière de Goyave and Sarcelle) are hydromorphic marine and continental soils, while Manche à eau has only hydromorphic marine soils (IRD, 2021). Continental soils are characterized by silt and clay with a compact structure, while hydromorphic marine salt soils are thin, composed of clay, sand, pebbles, and shell beds. Dequiedt et al. (2009) demonstrated that soil type greatly influences the structure of bacterial communities. Soil type could therefore impact the bacterial populations described in this study.

Predictive functional analysis performed using FAPROTAX revealed several predicted metabolic functions, including sulfate respiration, denitrification, sulfur respiration, nitrification, and the degradation of aromatic compounds. Despite the presence of these predicted functions, their relative abundances showed no marked differences between sites. This result can be explained by functional redundancy within the microbial communities at the three sites studied, i.e., the ability of different species to perform the same ecological functions (Louca et al., 2018). The potential presence of both denitrification and aromatic compound degradation functions in mangrove soils, as indicated by FAPROTAX, is consistent with the occurrence of anoxic conditions in these soils, as both processes are associated with low-oxygen environments. The presence of such conditions could therefore support the occurrence of other anaerobic degradation processes, including potential chlordecone degradation. Although its abundance does not differ significantly between sites, it is more strongly expressed in seaside areas. By way of comparison, the degradation of a polycyclic aromatic hydrocarbon by a *Marinobacter* strain has been observed in a coastal environment in southern China. This demonstrates that aromatic compound degradation can occur in coastal environments and suggests that similar degradation activities may also occur in mangrove ecosystems (Cui et al., 2016).

On the other hand, the presence of functional groups in the Manche à Eau is not surprising. The “aromatic compound degradation” function referenced in the FAPROTAX database is not specific to any particular molecule. Thus, the activity observed at the Manche à Eau could be linked to the degradation of other aromatic compounds present, such as Tris(2-chloroisopropyl)phosphate (Sebilo et al., 2025).

At the same time, analysis of the metabolic pathways via KEGG of the dominant genera in each sample from the three sites revealed the presence of four bacterial genera of interest in the chlordecone degradation process. The genera *Burkholderia*, *Dehalogenimonas*, *Ilumatobacter*, and *Thermodesulfovibrio* possess a complete vitamin B_12_ biosynthesis pathway, identified via KEGG modules [M00122] and [M00123]. This molecule is an essential cofactor in several enzymatic reactions, particularly those involved in reductive dehalogenation (Guo and Chen, 2018). Among these enzymes, reductive dehalogenase, which is anaerobic-dependent, is the most studied in the degradation of organohalogen compounds (Payne et al., 2015). The analysis revealed the presence of two bacterial genera containing a reducing dehalogenase: *Dehalogenimonas* and *Desulfomonile*. Previous genomic analysis of the *Desulfomonile* genus demonstrated the presence of a rdh operon containing a gene homologous to the dehalogenase reductase enzyme called rdh (Hug et al., 2013). This suggests that this bacterium could potentially be involved in the degradation of organohalogen compounds. On the other hand, the genus *Dehalogenimonas* is particularly promising due to the presence of a reductive dehalogenase. This metabolism has already been studied in this genus, particularly for the dechlorination of 1,2-dichloroethane (1,2-DCA) to ethene under high saline conditions, thanks to the enzyme DdeA (Qiao et al., 2022). The joint presence of this enzyme and its cofactor (vitamin B_12_) in the genus *Dehalogenimonas* suggests a potential ecological role in the degradation of chlordecone. Although this genus is present at all sites, it clearly dominates at Sarcelle, on the coast. This distribution, highlighted by 16S rDNA metabarcoding analyses, suggests a correlation with the results of the FAPROTAX analysis, which indicates the presence of functional groups involved in the degradation of aromatic compounds.

Indeed, the results of chlordecone measurements revealed a concentration of approximately 100 µg/kg in agricultural soils located along the edges of mangrove areas. For comparison, Cabidoche et al. (2009) measured a concentration of approximately 37.4 mg/kg, which corresponds to a concentration approximately 300 times higher than that measured in agricultural soils bordering the mangrove (Cabidoche et al., 2009).

The persistence of chlordecone observed in mangrove soils could nevertheless be related, at least in part, to a decoupling between the presence of microbial potential and its effective expression. Indeed, the presence of microorganisms potentially capable of contributing to the transformation of organochlorine compounds does not necessarily imply that the corresponding metabolic pathways are actively expressed under the environmental conditions encountered. This hypothesis can be considered in light of the recent study of Sebilo et al. (2026), who showed, in mangrove soils, the coexistence of a microbial potential associated with nitrate reduction and a low effective expression of this function. These authors interpret this decoupling as being partly a consequence of the low availability of nitrate, leading to the concept of a “functional nitrogen bank” (Sebilo et al., 2026). By analogy, it can therefore be hypothesized that certain micro-organisms potentially involved in chlordecone transformation could be present in mangrove soils without environmental conditions allowing the optimal expression of the corresponding metabolic pathways. However, this hypothesis remains to be experimentally tested, particularly through a comparison of microbial functional potential, its expression, and the bioavailability of chlordecone across the different compartments investigated.

These results reinforce the hypothesis that mangrove soils have a natural purification potential that is still under-exploited. Bacteria such as *Desulfomonile* and *Dehalogenimonas* are of great interest for targeted bioremediation of chlordecone, combining the two possible degradation pathways proposed (Della-Negra et al., 2021; Lomheim et al., 2021). This hypothesis is further supported by recent microcosm experiments conducted on chlordecone-contaminated mangrove sediments, which investigated the capacity of indigenous anaerobic microbial communities to promote chlordecone biodegradation under mesophilic conditions (Umar Zungum et al., 2026).

## Notes

### Competing Interest Statement

The authors have declared no competing interest.

